# Computation of the mitochondrial age distribution along the axon length

**DOI:** 10.1101/2021.09.12.459928

**Authors:** Ivan A. Kuznetsov, Andrey V. Kuznetsov

## Abstract

We describe a compartmental model of mitochondrial transport in axons, which we apply to compute mitochondrial age at different distances from the soma. The model predicts that at the tip of an axon that has a length of 1 cm, the average mitochondrial age is approximately 22 hours. The mitochondria are youngest closest to the soma and their age scales approximately linearly with distance from the soma. To the best of the authors’ knowledge, this is the first attempt to predict the spatial distribution of mitochondrial age within an axon. A sensitivity study of the mean age of mitochondria to various model parameters is also presented.

## 1. Introduction

Mitochondria generate easily usable chemical power for cells. In addition to their direct involvement in cellular respiration, they are also involved in apoptosis, calcium buffering, and many other important biological functions (Fan et al. 2001; Picard and McEwen 2018). Abnormalities in mitochondrial transport occur in many neurological disorders (Zheng et al. 2019). Recent research implicates α-synuclein-induced disruptions of mitochondrial trafficking in the development of Parkinson’s disease (PD) (Clague and Rochin 2016; Shahmoradian et al. 2019). Since packed mitochondria membrane fragments are one of the components found in Lewy bodies (Shahmoradian et al. 2019), investigating the transport of mitochondria is important for understanding the fundamentals of PD, for example in regard to possible energy deficits in dopaminergic neurons due to disruptions in mitochondrial axonal transport (Prots et al. 2018).

There are three pools of mitochondria in axons: anterogradely transported, retrogradely transported, and stationary (Hollenbeck and Saxton 2005). Understanding how old the mitochondria in these pools are and how their age depends on the distance from the soma is critical, as it yields insight into how mitochondria are renewed in the axon and the potential for dysregulation of mitochondrial transport in various diseases. Estimating the age of mitochondria may be important, for example, in evaluating the possible role of oxidative damage in the development of various neurodegenerative diseases (Uttara et al. 2009).

Renewal of mitochondrial proteins involves mitochondrial fusion and fission (Misgeld and Schwarz 2017). Fusion of old and new mitochondria leads to an exchange of old and new soluble and membrane-bound components, such as mitochondrial proteins (Malka et al. 2005; Patel et al. 2013). It is believed that mitochondrial fission leads to the separation of mutated copies of the mitochondrial genome into a separate mitochondrion, which is then recycled by the autophagosome (Twig et al. 2008; Safiulina and Kaasik 2013). Fission also serves as a mechanism that prevents mitochondria from becoming excessively long, so longer mitochondria are more likely to enter fission than shorter ones (Cagalinec et al. 2013). If fission events are unopposed, they can lead to complete fragmentation of the mitochondrial network (Ahola et al. 2019). Mitochondrial fusion accelerates mitochondrial fission, which is necessary to maintain a constant average length of mitochondria. Fission is important for autophagy, the mechanism of mitochondria removal (Twig et al. 2008).

Mitochondria are moved anterogradely by kinesin (in particular, by kinesin-1) motors and retrogradely by cytoplasmic dynein (Melkov and Abdu 2018). The motion of individual mitochondria is not continuous but rather interrupted by frequent pauses. Mitochondria move with an average velocity of 0.4-0.8 μm/s (Narayanareddy et al. 2014).

For investigating the distribution of the mean age of mitochondria in various compartments surrounding demand sites, Ferree et al. (2013) used a photoconvertible construct, termed MitoTimer, that switches from green to red fluorescence over a period of hours due to slow oxidation as it ages (red corresponds to older mitochondria and green corresponds to younger mitochondria). MitoTimer exhibits green fluorescence for the first 8 hours. After 24 hours it shows a mixture of green and red fluorescence (producing a yellow color). After 48-72 hours MitoTimer exhibits red fluorescence (Ferree et al. 2013; Hernandez et al. 2013; Gottlieb and Stotland 2015; Wilson et al. 2019). Experiments in Ferree et al. (2013) reported that mitochondria residing in compartments that surround the more distal demand sites are older than those residing in compartments surrounding more proximal demand sites. The ratio of red/green fluorescence depends on the rate of mitochondria synthesis in the soma, the rate of mitochondrial transport in the axon, the rates of fusion and fission (which equilibrate the MitoTimer protein content in mitochondria and mitochondria age), and autophagy.

A computational study (Patel et al. 2013) investigated mitochondria health, which is a proxy for mitochondrial membrane potential. An initial population of 150 unfused mitochondria was simulated. Recent research (Agrawal and Koslover 2021), which simulated mitochondria health as a decaying component, developed models for two maintenance models of mitochondria: one that assumed the exchange of mitochondrial proteins between moving and stationary mitochondria due to mitochondrial fusion and fission, and the other that assumed occasional replacement of stationary mitochondria with new ones arriving from the soma.

## 2. Materials and models

### 2.1. Governing equations

We treat the axon as a series of discrete compartments (representing mitochondria demand sites) and the age as the time since the production of a mitochondrion. The model developed in this paper accounts for stationary, anterogradely, and retrogradely moving mitochondria. We simulated a terminal with *N* high energy demand sites (hereafter demand sites) that contain stationary mitochondria (Fig. 1). The model developed in this research accounts for *N* resident and *2N* transiting (anterograde and retrograde) kinetic states and contains 3*N* ordinary differential equations (ODEs) (we used *N*=10 in this research).

**Fig. 1.**
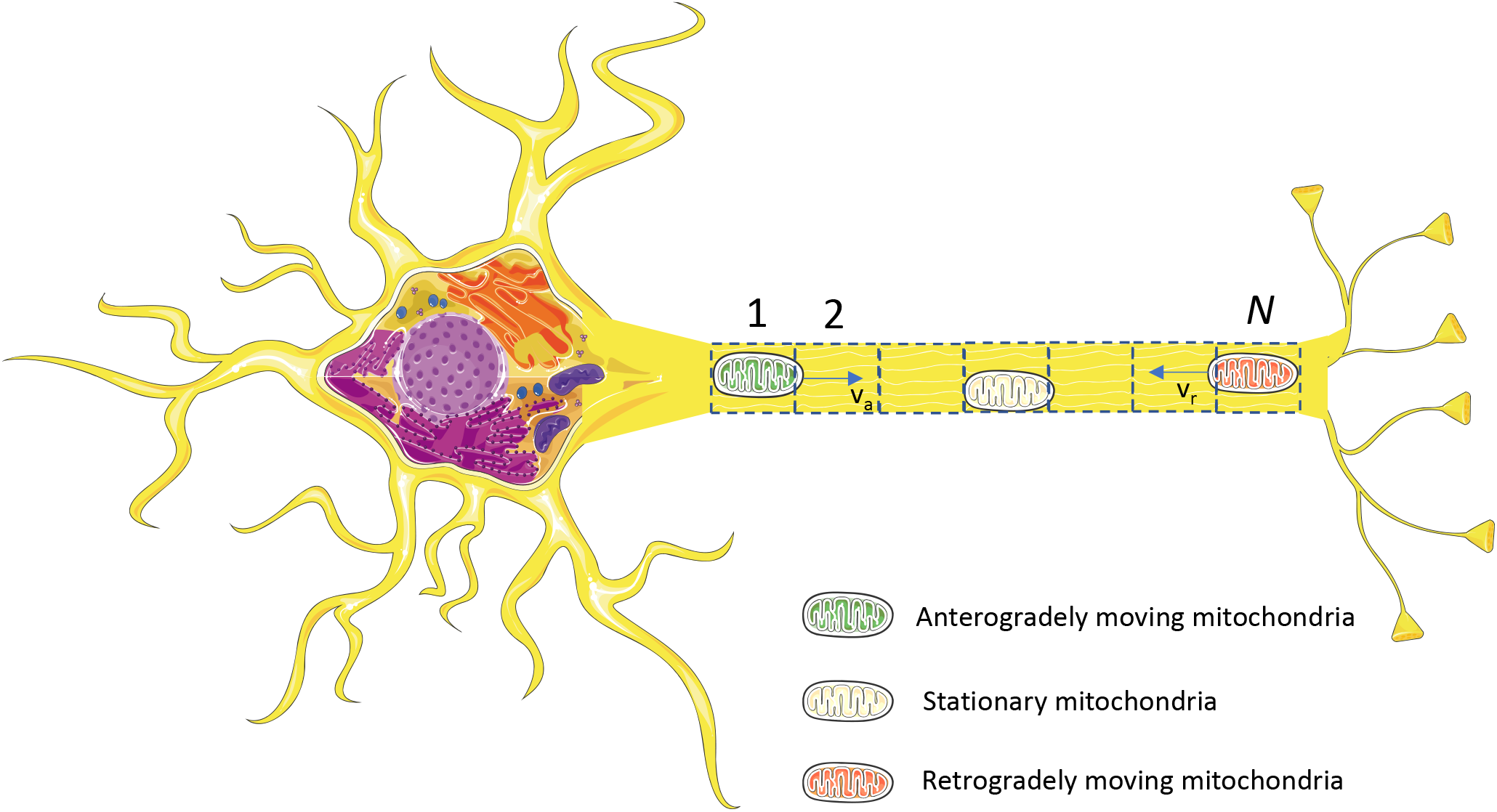
(a) A schematic diagram of a neuron with an axon which is assumed to contain *N* energy demand sites. Demand sites are numbered starting with the most proximal (site 1) to the most distal (site *N*) *Figure generated with the aid of servier medical art, licensed under a creative common attribution 3.0 generic license. http://smart.servier.com.*

We used a multi-compartment model (Anderson 1983; Jacquez 1985; Jacquez and Simon 1993) to formulate equations expressing the conservation of the total length of mitochondria in the compartments. Our rationale for representing the axon as a series of discrete compartments comes from our desire to use the methodology for computing the age distribution of cargos in compartmental systems developed in Metzler et al. (2018), Metzler and Sierra (2018). We used a similar compartmental approach to simulate the transport of dense core vesicles in *Drosophila* motoneurons (Kuznetsov and Kuznetsov 2019; Kuznetsov and Kuznetsov 2020). We numbered the demand sites in the axon terminal from the most proximal (site 1) to the most distal (site *N*) (Fig. 1). The demand sites have three compartments (representing resident, transiting-anterograde, and transiting-retrograde pools of mitochondria), see Fig. 2. To accurately model mitochondria fluxes between the demand sites, it is necessary to split the demand site into stationary, anterograde, and retrograde compartments. The ability of the model to predict mitochondria concentrations in these three different pools is also beneficial for biologists.

**Fig. 2.**
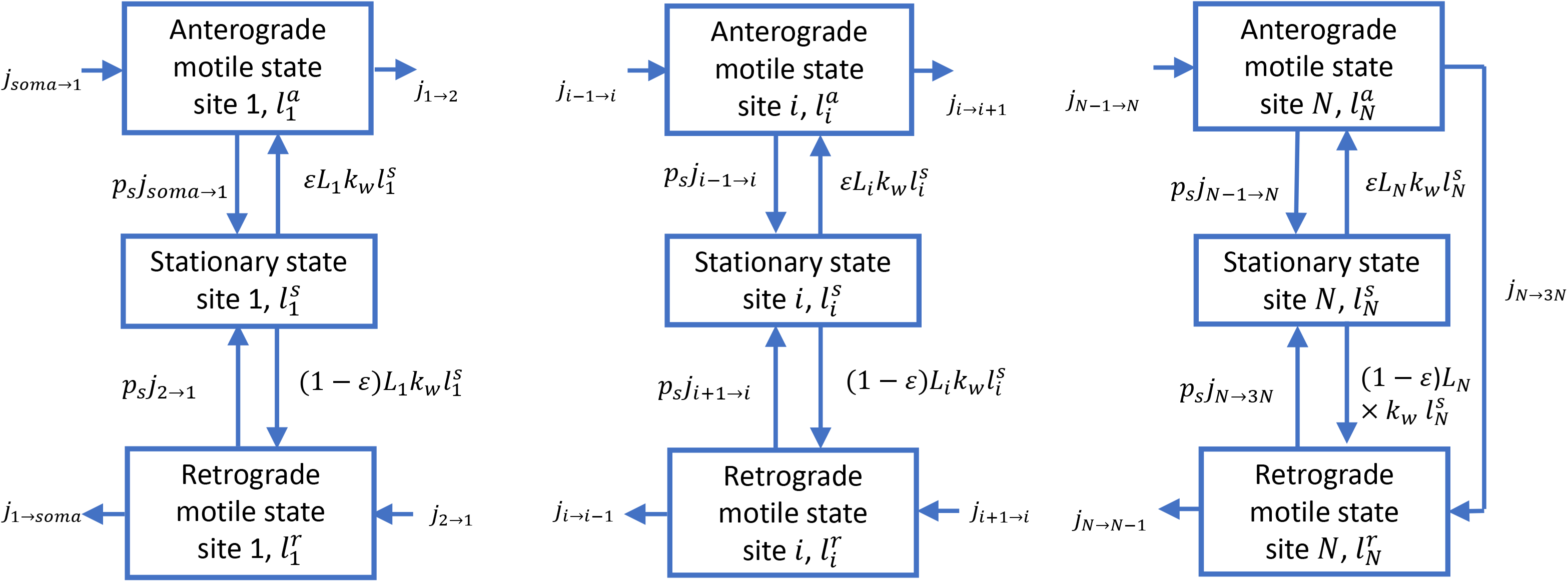
A diagram of a compartmental model showing transport in the compartments surrounding the demand sites in the axon. Arrows show mitochondria exchange between the transiting (anterograde and retrograde) states in adjacent demand sites, mitochondria capture into the stationary state and re-release from the stationary state. Mitochondria entering the axon are assumed to have zero age.

Since the length of an axon is much greater than its width, we characterized mitochondria concentration in a kinetic state by the total length of mitochondria in the kinetic state per unit axon length, measured in (μm of mitochondria)/μm. The total length of mitochondria is defined as the sum of the lengths of all mitochondria residing in the same kinetic state (stationary, anterograde, or retrograde) in the same compartment. The total length of mitochondria in the compartment for a particular kinetic state (Fig. 2) is the conserved quantity. The dependent variables utilized in the model are summarized in Table 1.

**Table 1.** Dependent variables in the model of mitochondrial transport and accumulation in the axon.

| Symbol | Definition | Units |
| --- | --- | --- |
| $l_i^s$ | Total length of stationary mitochondria per unit length of the axon in the compartment by the $i$th demand site | $(\mu\text{m of mitochondria})/\mu\text{m}$ |
| $l_i^a$ | Total length of anterogradely moving mitochondria per unit length of the axon in the compartment by the $i$th demand site | $(\mu\text{m of mitochondria})/\mu\text{m}$ |
| $l_i^r$ | Total length of retrogradely moving mitochondria per unit length of the axon in the compartment by the $i$th demand site | $(\mu\text{m of mitochondria})/\mu\text{m}$ |
| $j_{a \rightarrow b}$ | Flux of mitochondria length from state (compartment) $a$ to state (compartment) $b$ | ($\mu\text{m}$ of mitochondria) $\text{s}^{-1}$ |

First, the conservation of the total mitochondria length in the compartment around the most proximal demand site (site 1) is formulated. Stating this for the stationary pool of mitochondria gives the following equation (Fig. 2):

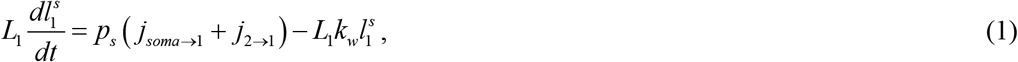

where *L_1_* is the length of an axonal compartment around the most proximal mitochondria demand site and *t* is the time. We assumed that the axonal length *L_ax_* is split into *N* compartments of equal length, for example, *L_1_ = L_ax_ / N*. The flux of mitochondria between the compartments is denoted by *j* (Table 1). Model parameters are defined in Table 2. For example, *p_s_* is the probability of mitochondria transitioning from a moving to a stationary state, *k_w_* is the kinetic constant characterizing the rate of mitochondria re-release from the stationary state into the motile states, and *j_soma→1_* is the flux at which mitochondria enter the axon.

**Table 2.** Parameters characterizing mitochondrial transport and accumulation in the axon.

| Symbol | Definition | Units | Value or range | Reference(s) | Value used in computations |
| --- | --- | --- | --- | --- | --- |
| $v_a$ | Average velocity of anterogradely moving mitochondria | $\mu\text{m s}^{-1}$ | 0.5 | Agrawal and Koslover (2021) | 0.5 |
| $v_r$ | Average velocity of retrogradely moving mitochondria | $\mu\text{m s}^{-1}$ | 0.5 | Agrawal and Koslover (2021) | 0.5 |
| $j_{soma \rightarrow 1}$ | Rate at which mitochondria enter the axon from the soma | mitochondria $\text{s}^{-1}$ | 0.0375 | Agrawal and Koslover (2021) | 0.0375<sup>a</sup> |
| $L_{ax}$ | Length of the axon | $\mu\text{m}$ | $10^4 - 10^5$ | Agrawal and Koslover (2021) | $10^4$ |
| $k_w$ | Kinetic constant characterizing the rate of mitochondria reentry into motile states (anterograde and retrograde) | $\text{s}^{-1}$ | $2 \times 10^{-4} - 2 \times 10^{-3}$<sup>b</sup> | Agrawal and Koslover (2021) | $2 \times 10^{-4}$,<br>$5 \times 10^{-4}$ |
| $\varepsilon$ | Portion of mitochondria leaving the stationary state that join the anterograde pool; $(1 - \varepsilon)$ join the retrograde pool | | 0.5 | Agrawal and Koslover (2021) | 0.5 |
| $f_s$ | Fraction of stationary mitochondria | | 0.5-0.6 | Agrawal and Koslover (2021) | 0.5 |
| $p_s$ | Probability that a motile mitochondrion would transition to a stationary state at a demand site | | $0.4 - 0.5$<sup>c</sup> | Agrawal and Koslover (2021) | 0.04, 0.4 |
| $N$ | Number of demand sites in the axon | | $5 - 100$<sup>d</sup> | Agrawal and Koslover (2021) | 10 |

<sup>a</sup> We estimated $j_{soma \rightarrow 1}$ using the formula suggested in Agrawal and Koslover (2021),
$Mv / (2L_{ax})$ , where $M$ is the number of mitochondria in the axon and $v$ is the average velocity of mitochondria motion in the axon. We used 1,500 mitochondria for $M$ , $0.5 \mu\text{m/s}$ for $v$ , and $1 \text{ cm}$ for $L_{ax}$ .
<sup>b</sup> We used Eq. (4) from Agrawal and Koslover (2021), $f_s = \frac{Nvp_s}{L_{ax}k_w + Nvp_s}$ . By solving this equation for $k_w$ using values given in Table 2 (for $N$ we used the range 10-100), we obtained that $k_w$ is in the range $2 \times 10^{-4}$ to $2 \times 10^{-3} \text{ s}^{-1}$ .
<sup>c</sup> We used $p_s = \frac{N_s}{2N}$ , where $N_s$ is an average number of stops that a mitochondrion makes during a back and forth journey in the axon (Agrawal and Koslover 2021).
<sup>d</sup> Number of demand sites in a real axon can be much larger than that. Ideally, the size of a compartment should represent a size of a mitochondria demand site (and the compartments should not be of the same size), but this would result in a very large number of compartments.

The conservation of mitochondria length in the anterograde pool of mitochondria around demand site 1 gives the following equation (Fig. 2):

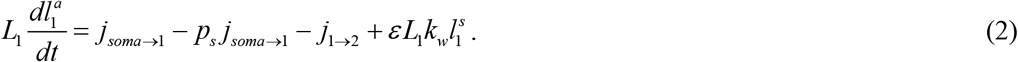

Unlike Agrawal and Koslover (2021), we did not assume that the mitochondria that reenter the motile state are split evenly between the anterograde and retrograde components. Instead, the portion of mitochondria that reenter the anterograde component is given by *ε* and the portion of mitochondria that reenter the retrograde component is given by (1-*ε*), see Fig. 2 and Table 2.

The conservation of mitochondria length in the retrograde pool of mitochondria around the most proximal demand site gives the following equation (Fig. 2):

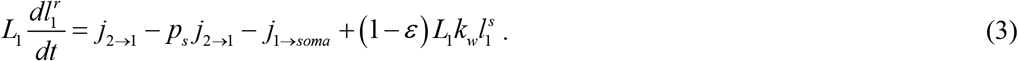

Next, the conservation of mitochondria length in the compartment around the *i*th demand site (*i*=2,…,*N*-1) is formulated. The conservation of mitochondria length in the stationary pool around the *i*th demand site gives the following equation (Fig. 2):

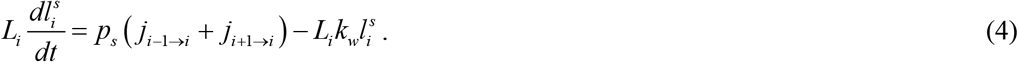

The conservation of mitochondria length in the anterograde pool in the *i*th demand site gives the following equation (Fig. 2):

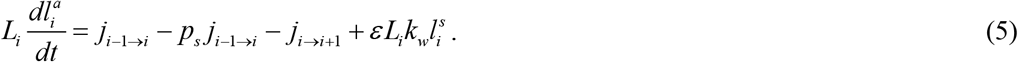

The conservation of mitochondria length in the retrograde pool in the *i*th demand site gives the following equation (Fig. 2):

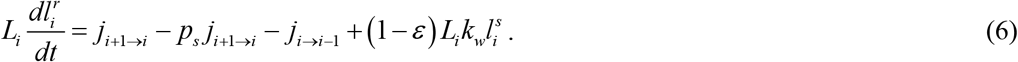

Finally, the conservation of mitochondria length in the compartment around the most distal demand site (site *N*) is formulated. As suggested by Agrawal and Koslover (2021), mitochondria that reach the distal end of the axon instantaneously switch from anterograde motors (kinesin) to retrograde motors (dynein) and continue to move retrogradely in the axon. This is simulated by *j_N→3N_* flux in Fig. 2.

The conservation of mitochondria length in the stationary pool gives the following equation (Fig. 2):

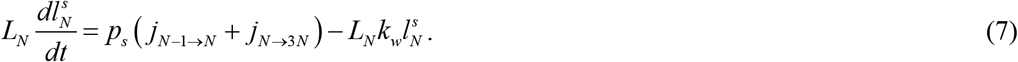

The conservation of mitochondria length in the anterograde pool gives the following equation (Fig. 2):

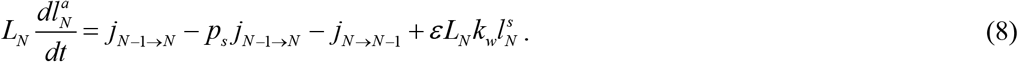

The conservation of mitochondria length in the retrograde pool gives the following equation (Fig. 2):

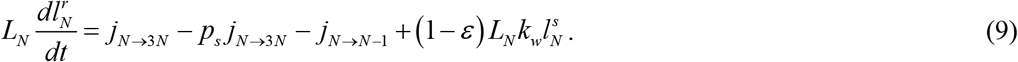

Eqs. (1)–(9) must be supplemented by the following equations for the mitochondria fluxes between the demand sites. *j_soma→1_* is an input model parameter; equations for other anterograde fluxes (Fig. 2) are

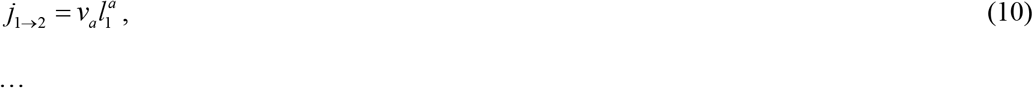

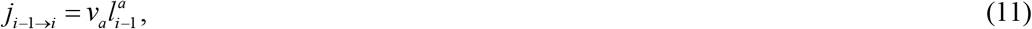

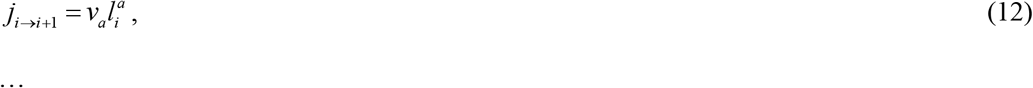

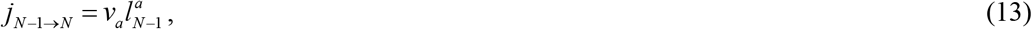

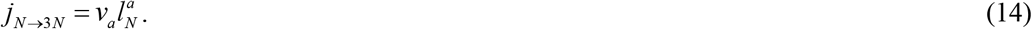

Equations for retrograde fluxes (Fig. 2) are

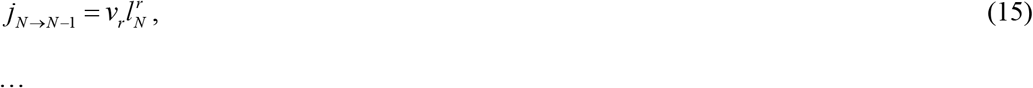

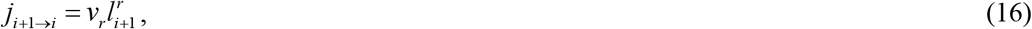

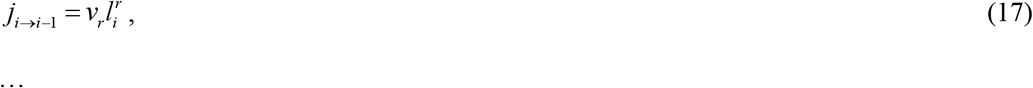

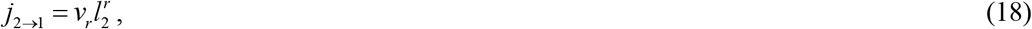

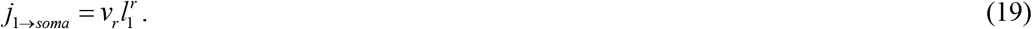

The model given by Eqs. (1)–(19) can also be replaced by a single set (valid between *x*=0 and *x*=*L_ax_*) of three partial differential equations (PDEs) (for anterograde, retrograde, and stationary mitochondria):

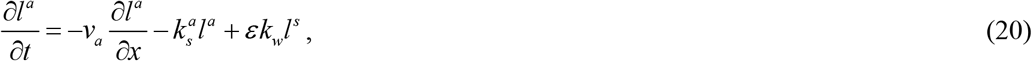

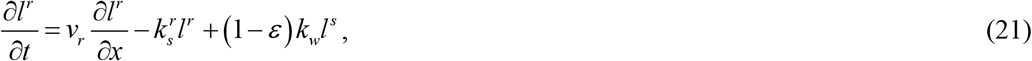

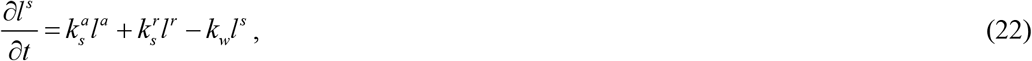

where

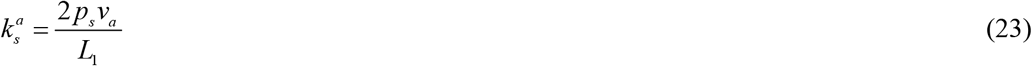

and

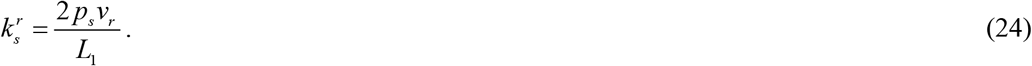

Eqs. (21)–(22) must be solved with the following boundary conditions:

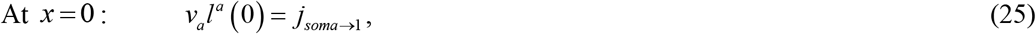

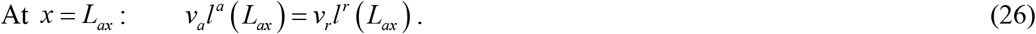

However, there is not a simple way to investigate how mitochondrial age varies throughout the axon utilizing PDEs (20)-(22). In order to more efficiently conduct our investigation, we resort to using discrete compartments (Fig. 2).

Since we seek the steady-state solution (which is of physical significance), the initial condition does not matter. We solved Eqs. (1)–(9) with zero initial conditions in all compartments and proceeded until the solution reached steady-state. The solution procedure involves solving initial value problems for three sets of ODEs. The numerical solution was obtained using MATLAB’s solver, ODE45 (MATLAB R2019a, MathWorks, Natick, MA, USA). The steady-state solution could also be obtained by setting the left-hand side of Eqs. (1)–(9) to zero and solving the obtained linear system for the densities in each compartment.

### 2.2. Model of mean age of mitochondria in the demand sites

To compute the age distributions of mitochondria in the stationary and moving states in the compartments surrounding the demand sites, the method described by Metzler et al. (2018), Metzler and Sierra (2018) was utilized. In order to use the methodology described in the above papers, Eqs. (1)–(9) were restated as the following matrix equation:

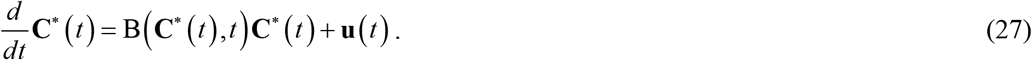

In Eq. (27) **C**\*, using the terminology of Metzler et al. (2018), is the state vector, which is defined by its components (**C**\* is a column vector):

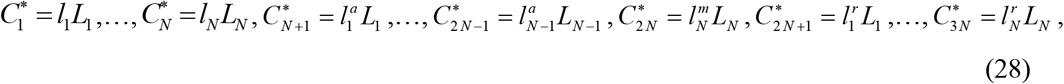

where

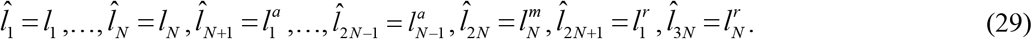

Rescaling the equations in the form given by Eqs. (27) and (28) is necessary because the conserved quantity is the total mitochondrial length in a certain compartment rather than its linear density.

The total length of mitochondria in stationary states is represented by the first *N* components of the state vector, the total length of mitochondria in anterograde states is represented by the next *N* components (*N*+1,…,2*N*), and the total length of mitochondria in retrograde states is represented by the last *N* components (2*N*+1,…,3*N*), see Fig. 2. Vector **u**(*t*) is defined below by Eqs. (60) and (61).

In our model, matrix B(3*N*,3*N*) is as follows. We accounted for the internal mitochondria fluxes between compartments, the external mitochondria flux entering the terminal from the axon, and the mitochondria flux leaving the terminal to move back to the axon (Fig. 2). Equations for the following elements of matrix B were obtained by analyzing mitochondria fluxes to and from the compartments surrounding the most proximal demand site:

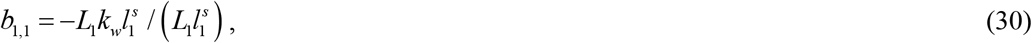

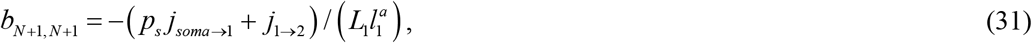

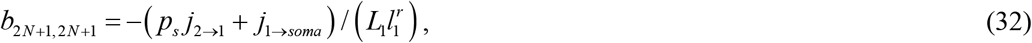

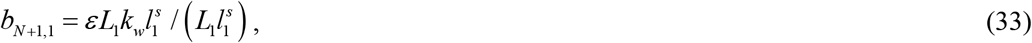

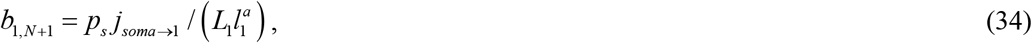

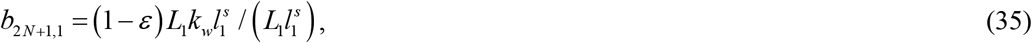

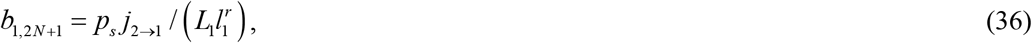

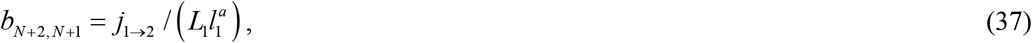

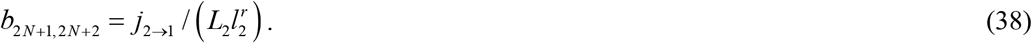

The analysis of mitochondria fluxes to and from the compartments surrounding demand site *i* (*i*=2,…,*N*-1) gives equations for the following elements of matrix B:

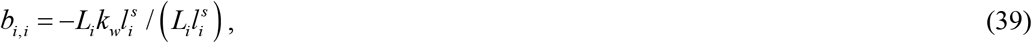

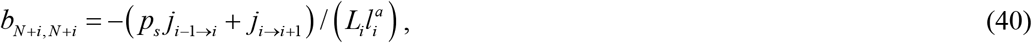

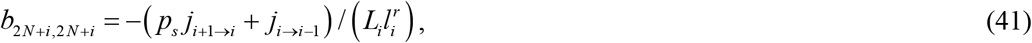

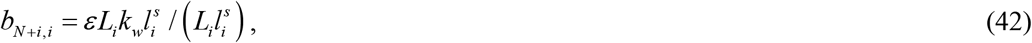

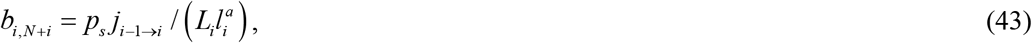

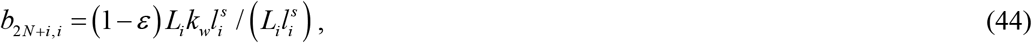

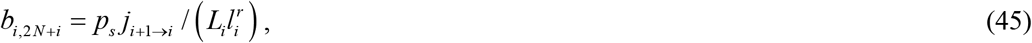

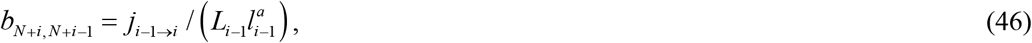

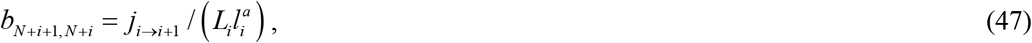

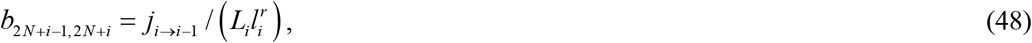

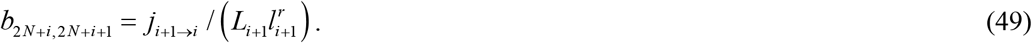

By analyzing mitochondria fluxes to and from the compartments surrounding the most distal demand site, equations for the following elements of matrix B were obtained:

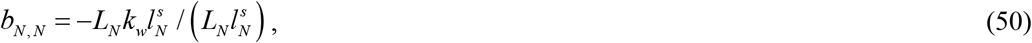

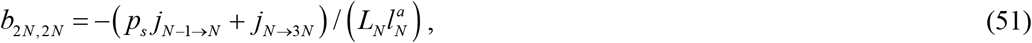

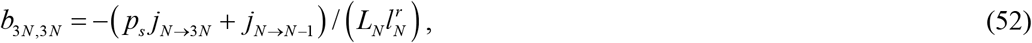

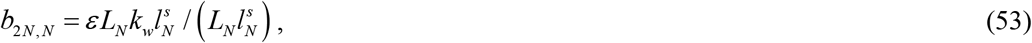

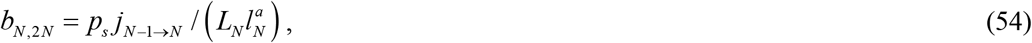

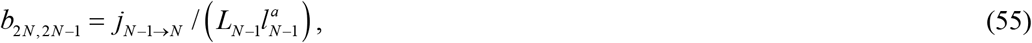

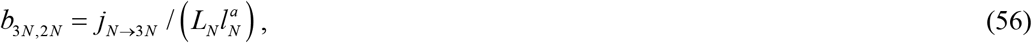

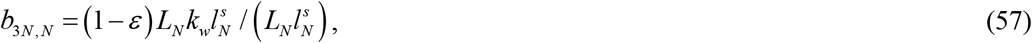

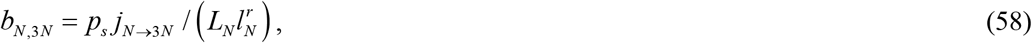

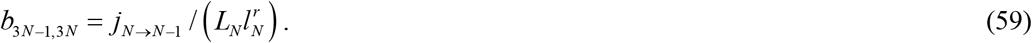

All other elements of matrix B, except for those given by Eqs. (30)–(59), are equal to zero. To better understand the physical significance of the elements of matrix B, consider a mitochondrion that enters the axon. According to our model, the behavior of this mitochondrion does not depend on the behavior of all other mitochondria. Then the elements on the main diagonal of matrix B determine how much time the mitochondrion will spend in a certain compartment, while the off-diagonal elements will determine how likely it is to transition to other compartments (Metzler and Sierra 2018).

The anterograde flux from the axon to the compartment with anterograde mitochondria by the most proximal demand site, *j_soma→1_*, is the only flux entering the terminal. Our model assumes that all mitochondria that leave the terminal (their flux is *j_soma→1_*) return to the soma for degradation, and none reenter the terminal. We also assumed that mitochondria that enter the terminal (their flux is *j_soma→1_*) are all newly synthesized in the soma and their age at the time of entry is equal to zero. This means that our model does not account for the time it takes mitochondria to travel from the soma to the terminal. The mitochondrial age calculated here should be understood as the time since mitochondria entered the terminal.

The following equation yields the *N*+1^th^ element of column vector **u**:

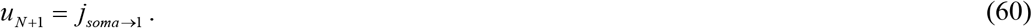

Because no other external fluxes enter the terminal,

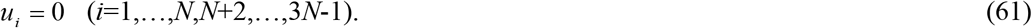

The state transition matrix Φ, which can also be viewed as a matrix-valued Green’s function, is defined in Metzler et al. (2018). It may be determined by solving the following matrix equation:

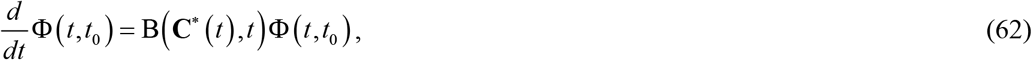

where *t*_0_ is the initial time. In our case *t*_0_ = 0.

The initial condition for Eq. (62) is

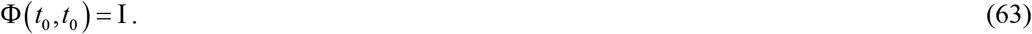

In Eq. (63) symbol I denotes an identity matrix.

We assumed that all mitochondria that enter the terminal are new. The age density of mitochondria entering the terminal after *t =* 0 can then be determined from the following equation:

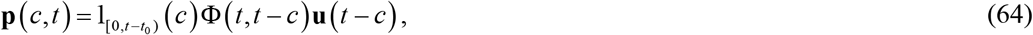

where 1_[0 t-t_o_)_ is the indicator function, which equals 1 if 0 ≤*c* < *t-t_0_* and equals 0 otherwise. The age density of mitochondria is the ratio of the total length of mitochondria in the compartment which have an age between *T* and *T+dT* over the duration of the interval *dT*. An integral of the age density of mitochondria over any time period with respect to time gives the length of mitochondria having an age within that time range.

The mean age of mitochondria in demand sites is (Rasmussen et al. 2016):

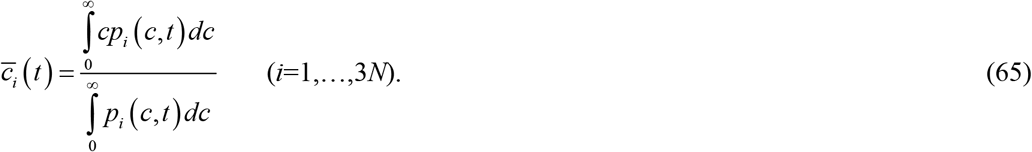

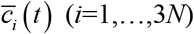 may be found by solving the following mean age system (Metzler et al. 2018; Rasmussen et al. 2016):

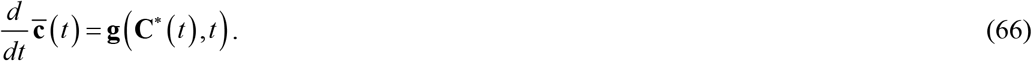

The initial condition for Eq. (66) is

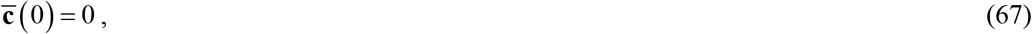

where 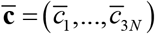. Vector **g** is defined as follows:

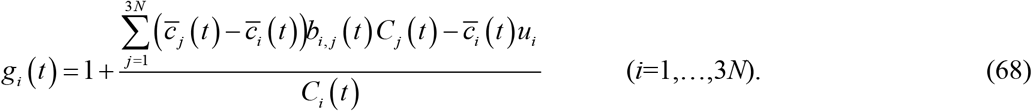

The following vectors were then defined to represent the mean age of mitochondria in the stationary, anterograde, and retrograde states in the demand sites, respectively:

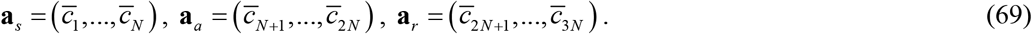

The arrays **a**_*s*_, **a**_*a*_, and **a**_*r*_ are one-dimensional and of size *N*.

Numerical solutions were again obtained utilizing MATLAB’s solver, ODE45. The error tolerance parameters, RelTol and AbsTol, were set to 10 ^-6^ and 10 ^-8^, respectively. We checked that lowering RelTol and AbsTol did not influence the solutions. The results were also checked using the formulas reported in Metzler and Sierra (2018). In particular, the solution of Eq. (27) for a steadystate can be obtained as:

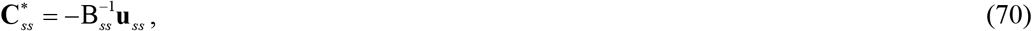

where the superscript–1 on a matrix denotes the inverse of the matrix and the subscript *ss* denotes steady-state. The mean ages of mitochondria in various compartments displayed in Fig. 2 at steady-state can be obtained as:

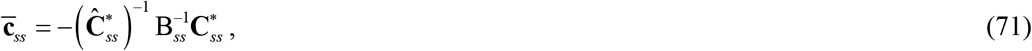

where

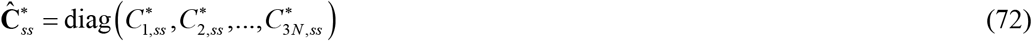

is the diagonal matrix with the components of the state vector, defined in Eq. (28), on the main diagonal. These components (*C*_1,ss_*, *C*_2,ss_*, etc.) are calculated at steady-state. Also, in Eq. (71)

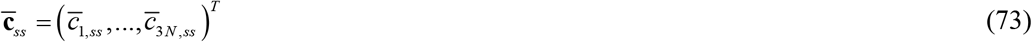

is the column vector composed of the mean ages of mitochondria in various compartments (superscript *T* means transposed).

## 3. Results

### 3.1. Estimation of the average age of mitochondria in the most distal demand site

According to Fig. 4C in Ferree et al. (2013), at some point in time the green/red fluorescence ratio, *y*, linearly decreases with an increase in the distance from the nucleus as:

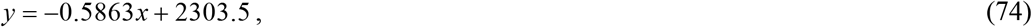

where *x* is in μm and *y* is in arbitrary units. From Eq. (74) it follows that

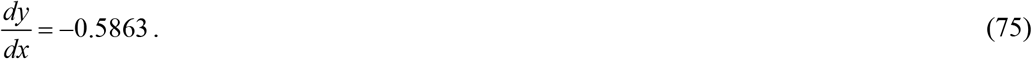

In Ferree et al. (2013) MitoTimer production was induced by exposure to doxycycline. Green fluorescence starts approximately 5 hours after doxycycline induction. At that point in time, red fluorescence is equal to zero (Fig. 2A in Ferree et al. 2013). According to Fig. 2A in Ferree et al. (2013), the red/green fluorescence ratio, which is the reciprocal of *y*, is approximately given by

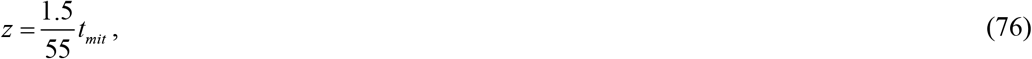

where *t_mit_* must be given in hours (*t_mit_ =* 0 corresponds to approximately 5 hours after doxycycline induction, the time when green fluorescence starts and red fluorescence is equal to zero, see Fig. 1C from Ferree et al. 2013).

At *x* =0 we assumed that all mitochondria are newly synthesized and thus their age is zero. For the estimate of the mitochondrial age at the axon terminal, we assumed a direct relation between the age of a mitochondrion, *t_mit_*, and its distance from the soma (this assumption is supported by Shlevkov and Schwarz 2017):

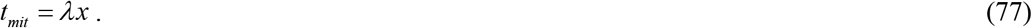

Combining Eqs. (75), (76), and (77), it follows that

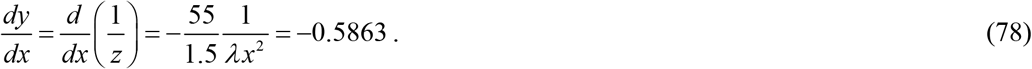

We used Eq. (78) at *x* = 200 μm (one-third of the range displayed in Fig. 4C of Ferree et al. 2013) and solved for *λ*. The result is

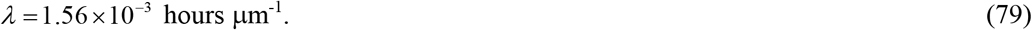

Calculating the age of mitochondria at the axon tip (at *x* = *L_ax_*) leads to

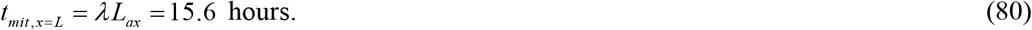

This is consistent with the expected half-life of mitochondrial proteins (days or weeks) (Misgeld and Schwarz 2017). It is interesting that data obtained in Mandal et al. (2018) using Timer fluorescent protein support rapid turnover of mitochondria in axon terminals and suggest that the residence time of the protein in the synapse is approximately 3 hours. Mandal et al. (2021) reported that complete turnover of mitochondria in axon terminals of zebrafish neurons occurs within 24 hours.

### 3.2. Total length of various populations of mitochondria per unit length of the axon and mean age of mitochondria in various demand sites

Most of the parameter values given in Table 2 are extracted from Agrawal and Koslover (2021) for the case corresponding to the optimal mitochondria health. *k_w_* = 5×10^-4^ s^-1^ was chosen as the base case because, as demonstrated later, the predicted mitochondrial age for this value roughly matches that observed in experimental data.

The compartmental model predicts spatially uniform densities of anterograde, retrograde, and stationary mitochondria throughout the axon (Fig. 3). The same density distributions are obtained by solving the PDE model (20)-(22) (Fig. 4), which confirms the equivalence of the two models. The result is expected because mitochondria should be distributed in an approximately uniform manner throughout the axon. This distribution allows the mitochondria satisfy the energy demand of all demand sites.

**Fig. 3.**
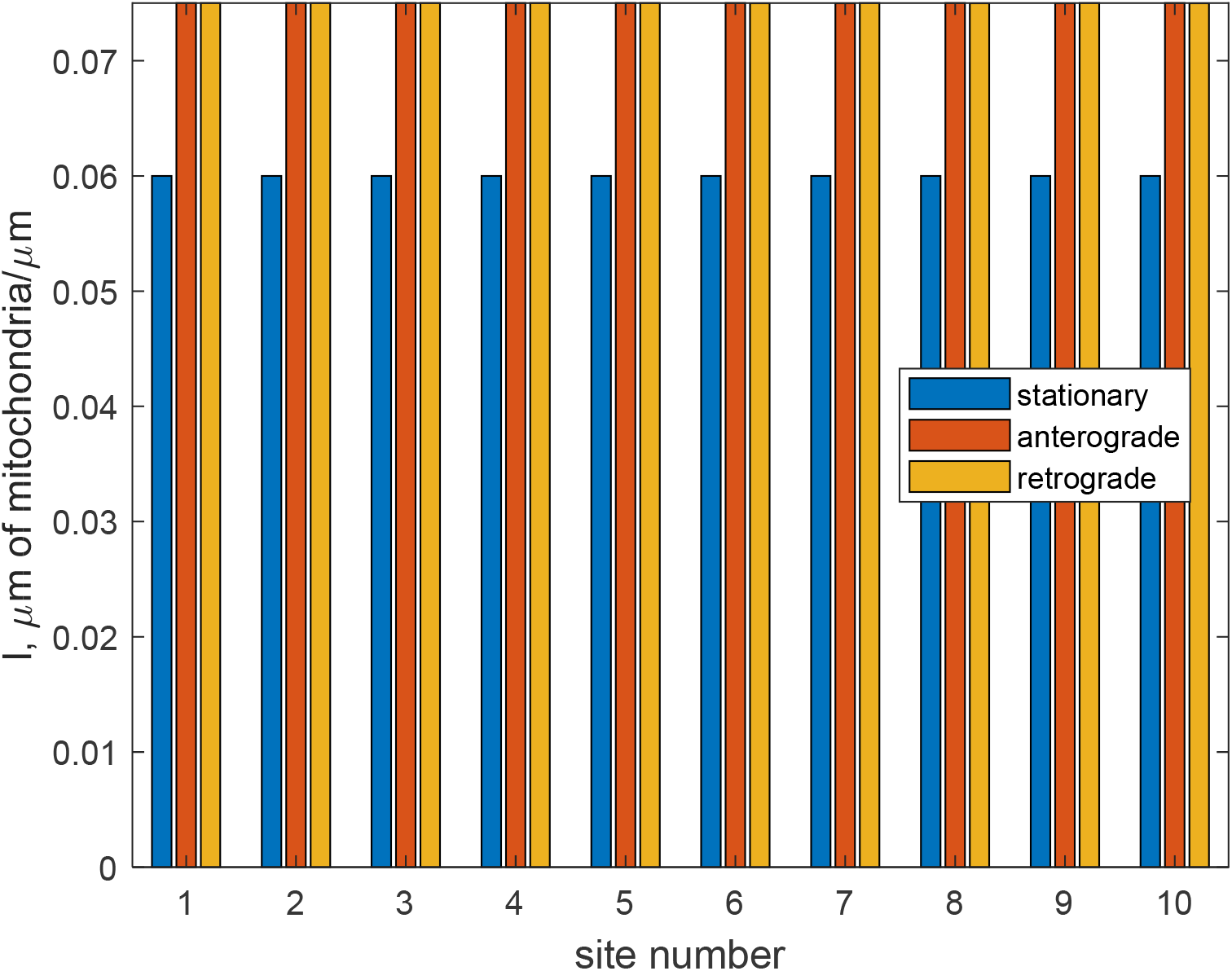
Steady-state values of the total length of stationary, anterogradely moving, and retrogradely moving mitochondria per unit length of the axon in the compartment by the *i*th demand site. *k_w_* = 5×10^-4^ s^-1^, *p_s_* = 0.4.

**Fig. 4.**
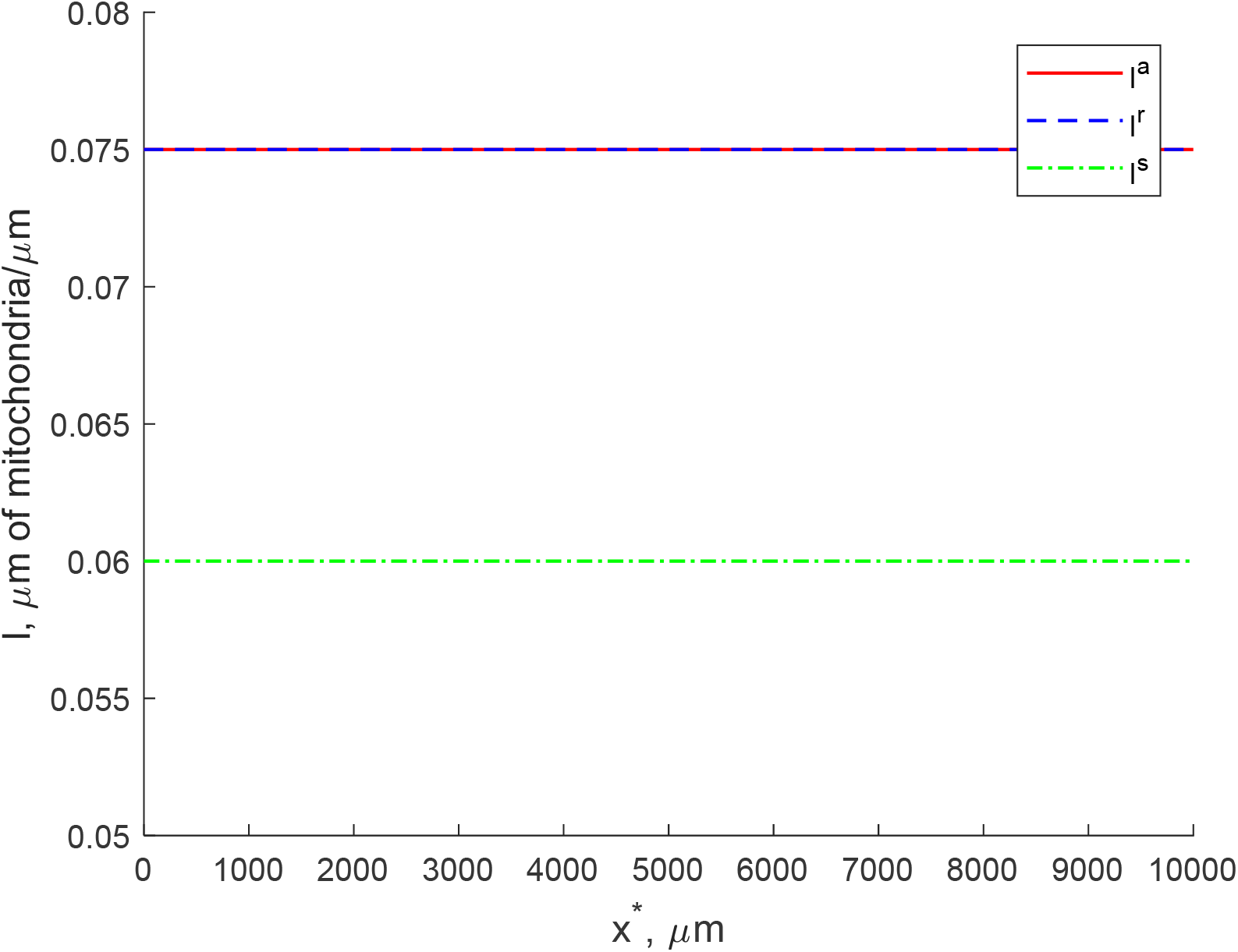
Steady-state values of the total length of stationary, anterogradely moving, and retrogradely moving mitochondria per unit length of the axon obtained by solving PDEs (20)-(22) with boundary conditions (25), (26). *k_w_* = 5×10^-4^ s^-1^, *p_s_* = 0.4.

Computations with *k_w_* = 5×10^-4^ s^-1^ (Table 2) yield the mean age of stationary mitochondria in demand site 10 (the most distal demand site) to be approximately 21.6 hours (Fig. 5), which is consistent with the estimate given by Eq. (80). It is known that older mitochondria may have impaired function due to accumulated oxidative damage (Saxton and Hollenbeck 2012). For short axons, oxidative damage is unlikely to be a contributing factor in mitochondrial diseases. For example, since the age of mitochondria at the tip of an axon with a length of 1 cm is only 22 hours, they will most likely not experience oxidative damage.

**Fig. 5.**
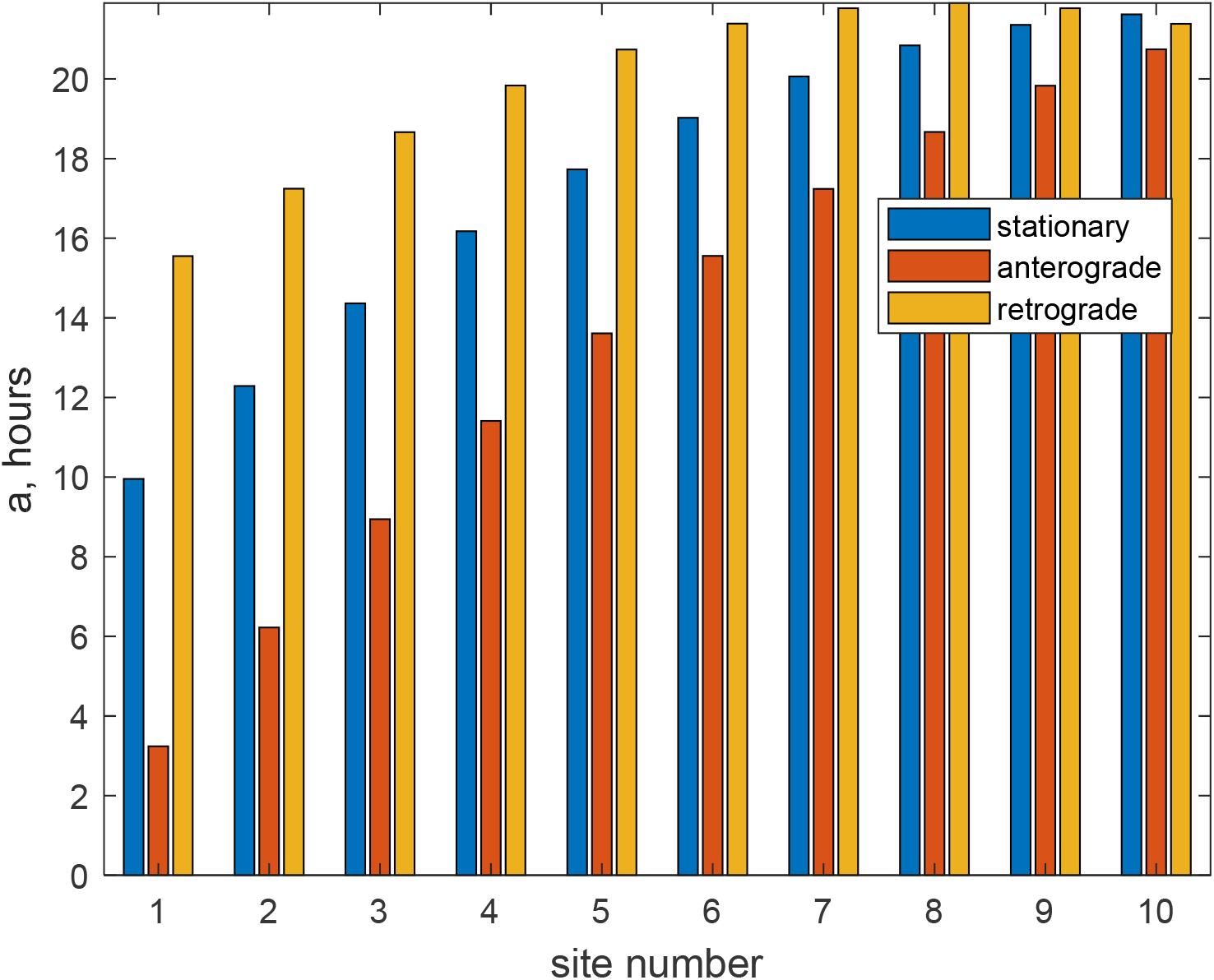
Steady-state values of the mean age of stationary, anterogradely moving, and retrogradely moving mitochondria in the compartment by the *i*th demand site. *k_w_* = 5×10^-4^ s^-1^, *p_s_* = 0.4.

The mean ages of stationary, anterograde, and retrograde mitochondria increase from the proximal to more distal demand sites. This is in agreement with results reported in Ferree et al. (2013), Shlevkov and Schwarz (2017), who found that the age of mitochondria increased approximately linearly with distance from the soma. As expected, the mean age of anterograde mitochondria is the smallest and the mean age of retrograde mitochondria is the largest.

The mean age of stationary mitochondria of 21.6 hours in the most distal demand site is severalfold greater than the time it takes for the mitochondria to travel from the soma to the most distal demand site without stopping:

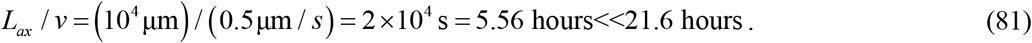

Some recent research (Smit-Rigter et al. 2016; Lees et al. 2020) suggest that *in vivo* mitochondria may stay in the stationary state for longer periods of time than *in vitro*. The model parameters can be adjusted to characterize transitions between anterograde, retrograde, and stationary states. This will affect the average time that mitochondria spend stopping near the demand sites in particular neural cells. Although the mean age of mitochondria depends on the values of model parameters, we believe that the important physics of mitochondria transport is captured by our model. Computations with *k_w_* = 2×10 ^-4^ s^-1^ (2.5 times less the value used in computing Fig. 5) and *p_s_* = 0.4 yield the mean age of stationary mitochondria in demand site 10 (the most distal demand site) to be approximately 31.5 hours (Fig. 6a).

**Fig. 6.**
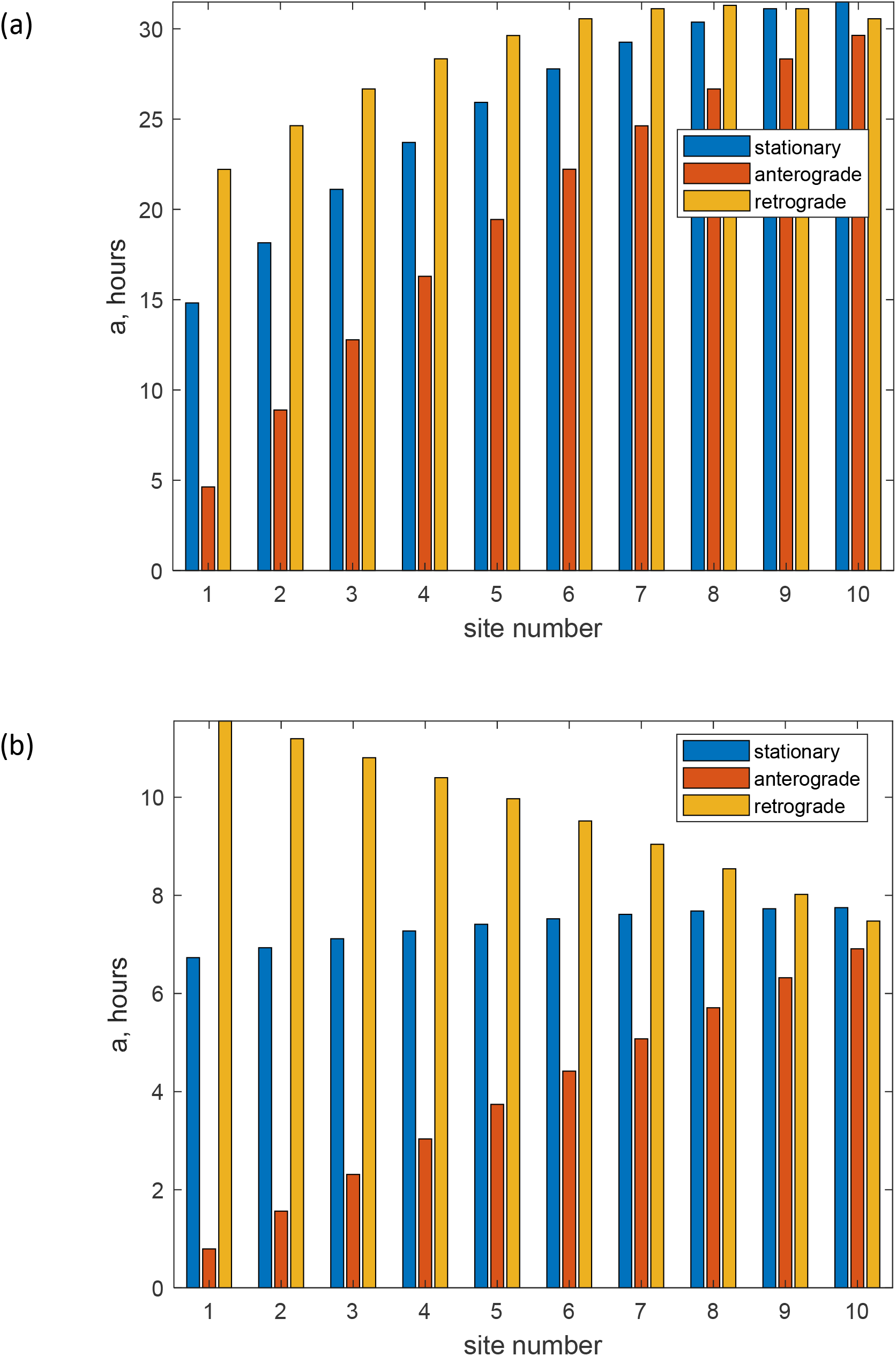
Steady-state values of the mean age of stationary, anterogradely moving, and retrogradely moving mitochondria in the compartment by the *i*th demand site for the case of (a) *k_w_* = 2 ×10^-4^ s^-1^, *p_s_* = 0.4, (b) *k_w_* = 5×10^-4^ s^-1^, *p_s_* = 0.04.

The smaller mean ages of retrograde mitochondria in proximal demand sites in Figs. 5 and 6a are explained by the capture of younger mitochondria from the anterograde state into the stationary state and then their subsequent re-release into the retrograde state (Fig. 2). If the value of parameter *p_s_* that characterizes the probability of capture of motile mitochondria into the stationary state is decreased, the age of retrograde mitochondria increases from the distal (site 10) to proximal (site 1) demand sites, as mitochondria get older as they are transported back to the soma (Fig. 6b).

### 3.3. Investigating sensitivity of the mean age of mitochondria in demand sites to model parameters

To investigate the sensitivity of the solution to model parameters, we calculated the local sensitivity coefficients. These are first-order partial derivatives of the observables with respect to model parameters (Beck and Arnold 1977; Zadeh and Montas 2010; Zi 2011; Kuznetsov and Kuznetsov 2019). For example, the sensitivity coefficient of the mean age of resident mitochondria in demand sites to parameter *k_w_* at steady-state (ss) can be calculated as follows:

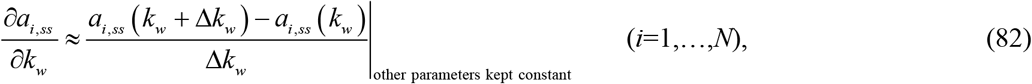

where Δ*k_w_* = 10^-1^ *k_w_* (the accuracy was tested by using various step sizes).

The sensitivity coefficients were non-dimensionalized by introducing relative sensitivity coefficients (Zadeh and Montas 2010; Kacser et al. 1995), defined as (for example):

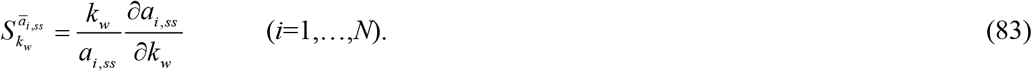

The dimensionless sensitivity of the mean age of mitochondria to various model parameters is displayed in Figs. 7–9 and Fig. S1. Out of four investigated parameters, the mean age is most sensitive to parameter *ε*, less sensitive to parameters *p_s_* and *v*, and least sensitive to parameter *k_w_*.

**Fig. 7.**
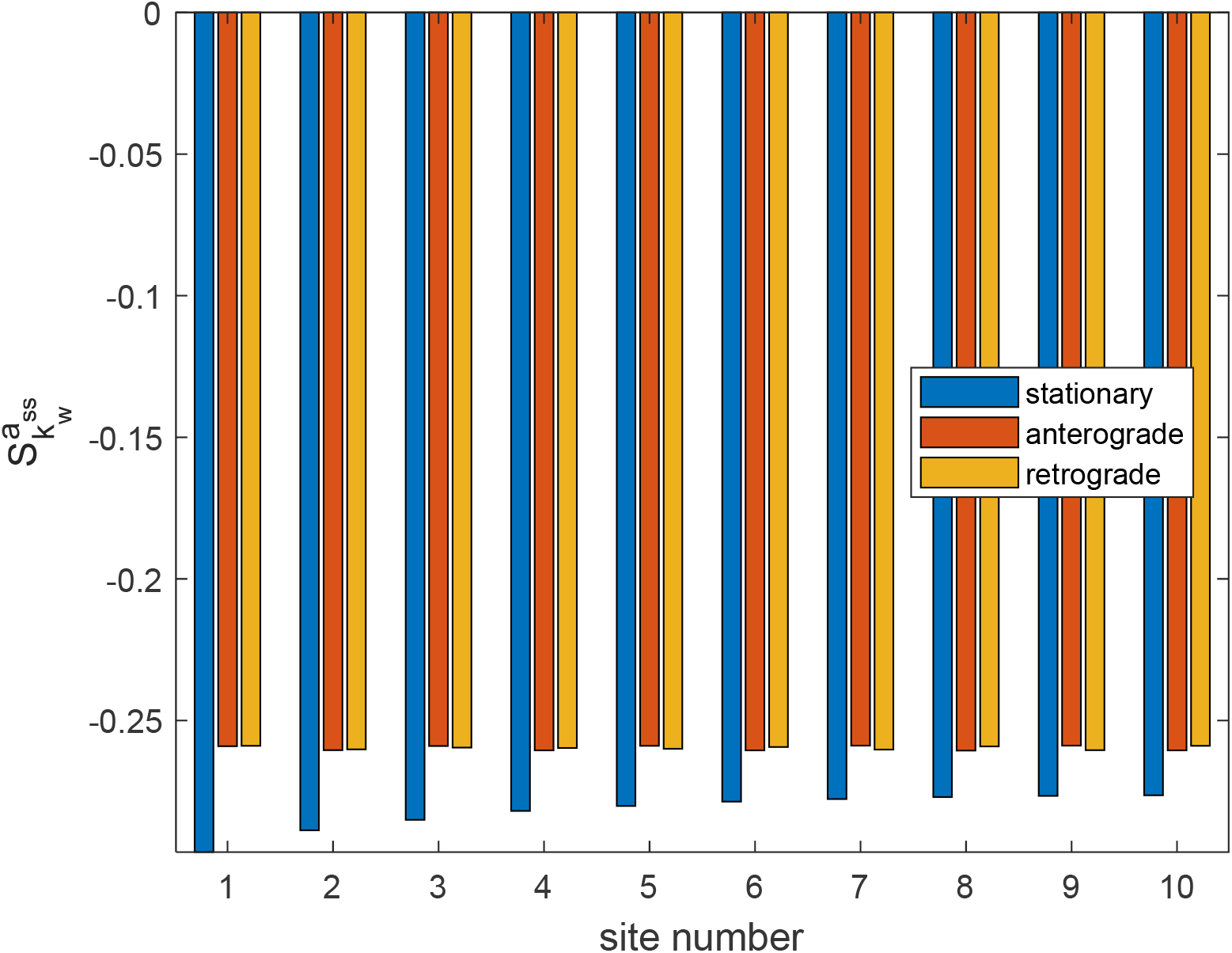
Dimensionless sensitivity of the mean age of mitochondria to the kinetic constant characterizing the rate of mitochondria reentry into motile states, *k_w_*, versus the site number. Sensitivity is analyzed around *k_w_* = 5×10^-4^ s^-1^, *p_s_* = 0.4. Computations were performed with Δ*k_w_* = 10^-1^*k_w_*. A close result was obtained for Δ*k_w_* = 10^-2^*k_w_*.

The dimensionless sensitivity of the mean age of mitochondria to the restating rate, *k_w_*, is negative in all demand sites (Fig. 7). This is because for the case with a larger restarting rate mitochondria spend less time in the stationary state. The negative sensitivity in Fig. 7 is consistent with the results in Figs. 5 and 6, which indicate that a reduction in *k_w_* increases the mean age of mitochondria. This is because fewer mitochondria (for smaller *k_w_*) escape from the stationary state and transition to the moving (anterograde or retrograde) states.

The dimensionless sensitivity of the mean age of mitochondria to the stopping probability, *p_s_*, is positive in all demand sites (Fig. 8). This is because larger values of *p_s_* correspond to more mitochondria transitioning to the stationary state.

**Fig. 8.**
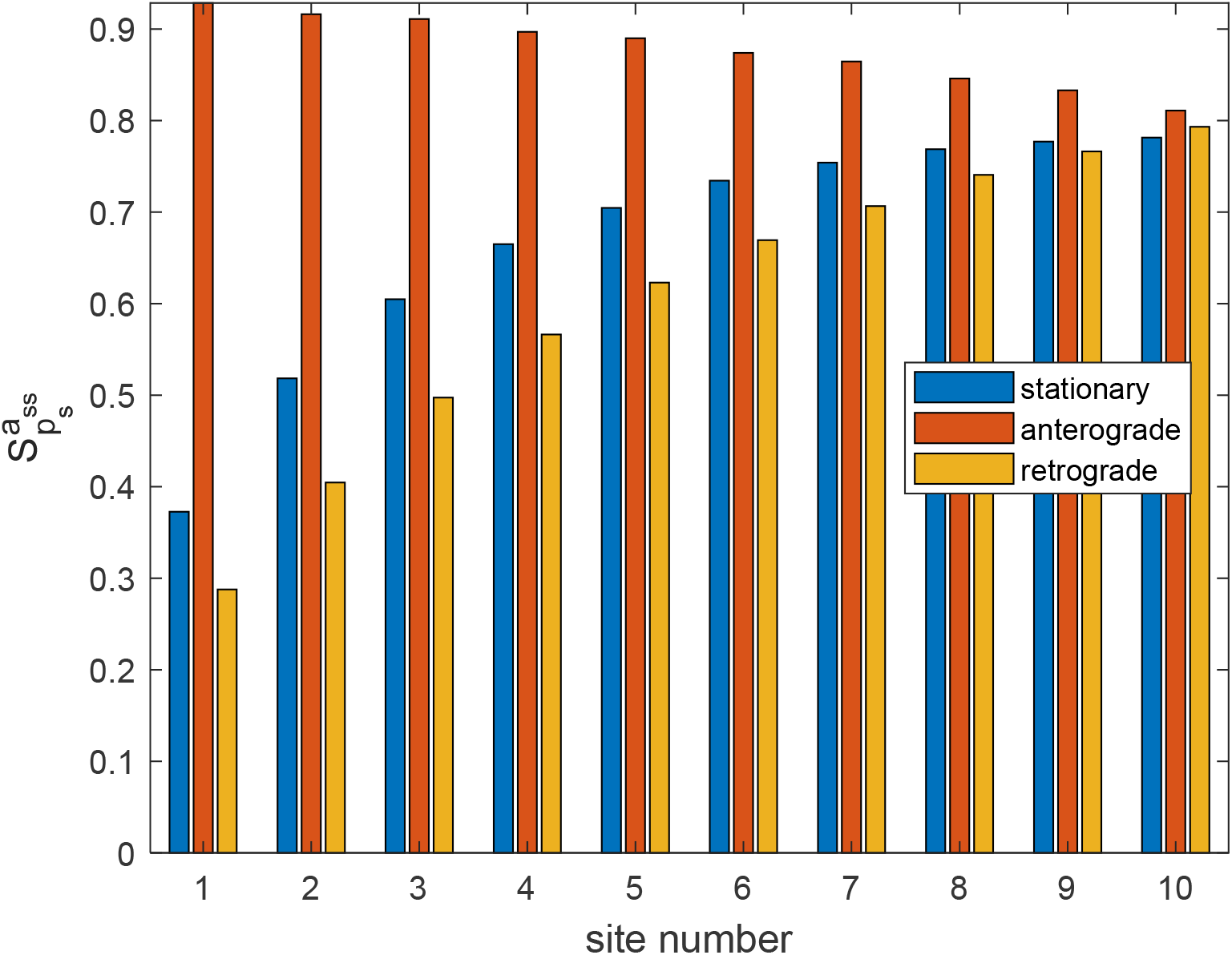
Dimensionless sensitivity of the mean age of mitochondria to the probability that a motile mitochondrion would transition to a stationary state at a demand site, *p_s_*, versus the site number. Sensitivity is analyzed for *k_w_* = 5×10^-4^ s^-1^, *p_s_* = 0.4. Computations were performed with Δ*p_s_* = 10^-1^*p_s_*. A close result was obtained for Δ*p_s_* = 10^-2^ *p_s_*.

The dimensionless sensitivity of the mean age of mitochondria to the anterograde bias, *ε*, is positive in all demand sites (Fig. 9). This is because mitochondria that transition to the anterograde state end up spending more time in the axon than mitochondria that transition to the retrograde state, which exit the axon sooner.

**Fig. 9.**
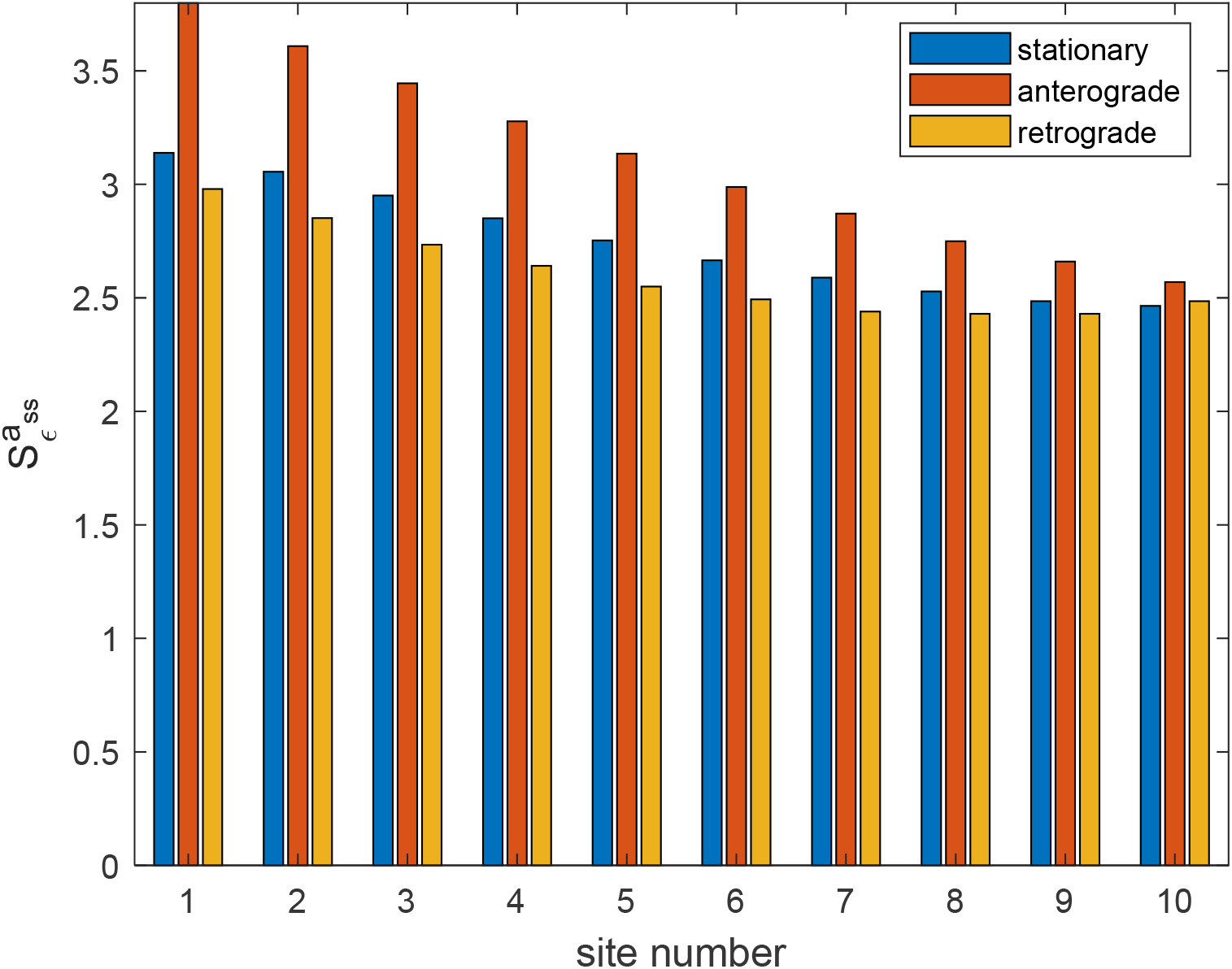
Dimensionless sensitivity of the mean age of mitochondria to the portion of mitochondria leaving the stationary state that join the anterograde pool, *ε*, versus the site number. Sensitivity is analyzed for *k_w_* = 5×10 ^4^ s^-1^, *p_s_* = 0.4. Computations were performed with △*ε* =10 ^-1^*ε*. A close result was obtained for Δ*ε* = 10^-2^ *ε*.

The dimensionless sensitivity of the mean age of mitochondria to mitochondria velocity, *v*, is negative in all demand sites (Fig. S1). This is because the faster movement of mitochondria corresponds to less time that mitochondria spend in the axon.

## 4. Discussion, limitations of the model and future directions

The model gives quantitative predictions for the distribution of mitochondrial mean age throughout the axon length that can be tested experimentally, indicating that the model has predictive value.

The model predicts that the mean age of stationary mitochondria in the most distal demand site is approximately 21.6 hours. This is severalfold greater than the time it takes for mitochondria to travel without stopping from the soma to the farthest demand location. To quantify the dependence of the solutions on model parameters, we employed analysis of the dimensionless sensitivity of the mean age of mitochondria at demand sites.

Future development of the model should address the fact that mitochondria that return to the soma from the axon are usually not destroyed, but rather fuse with a resident somatic pool of mitochondria (Misgeld and Schwarz 2017). Therefore, our assumption that mitochondria have zero age at the entrance to the axon is a simplifying one.

Loss/degeneration of mitochondria via mitophagy is an important aspect that needs to be incorporated into the model in future research. However, as Agrawal and Koslover (2021) pointed out, it is difficult to do this within the continuum approach because mitophagy involves selectively removing mitochondria with low membrane potential (in our approach these could be viewed as mitochondria that reached a certain age) by engulfing them in autophagosomes and then transporting them to the soma for degradation in lysosomes. This was done in Agrawal and Koslover (2021) by using a discrete stochastic simulation model.

## Acknowledgment

IAK acknowledges the fellowship support of the Paul and Daisy Soros Fellowship for New Americans and the NIHINational Institute of Mental Health (NIMH) Ruth L. Kirchstein NRSA (F30 MH122076-01). AVK acknowledges the support of the National Science Foundation (award CBET-2042834) and the Alexander von Humboldt Foundation through the Humboldt Research Award.

## Supplemental Materials

### S1. Supplementary figures

**Fig. S1.**
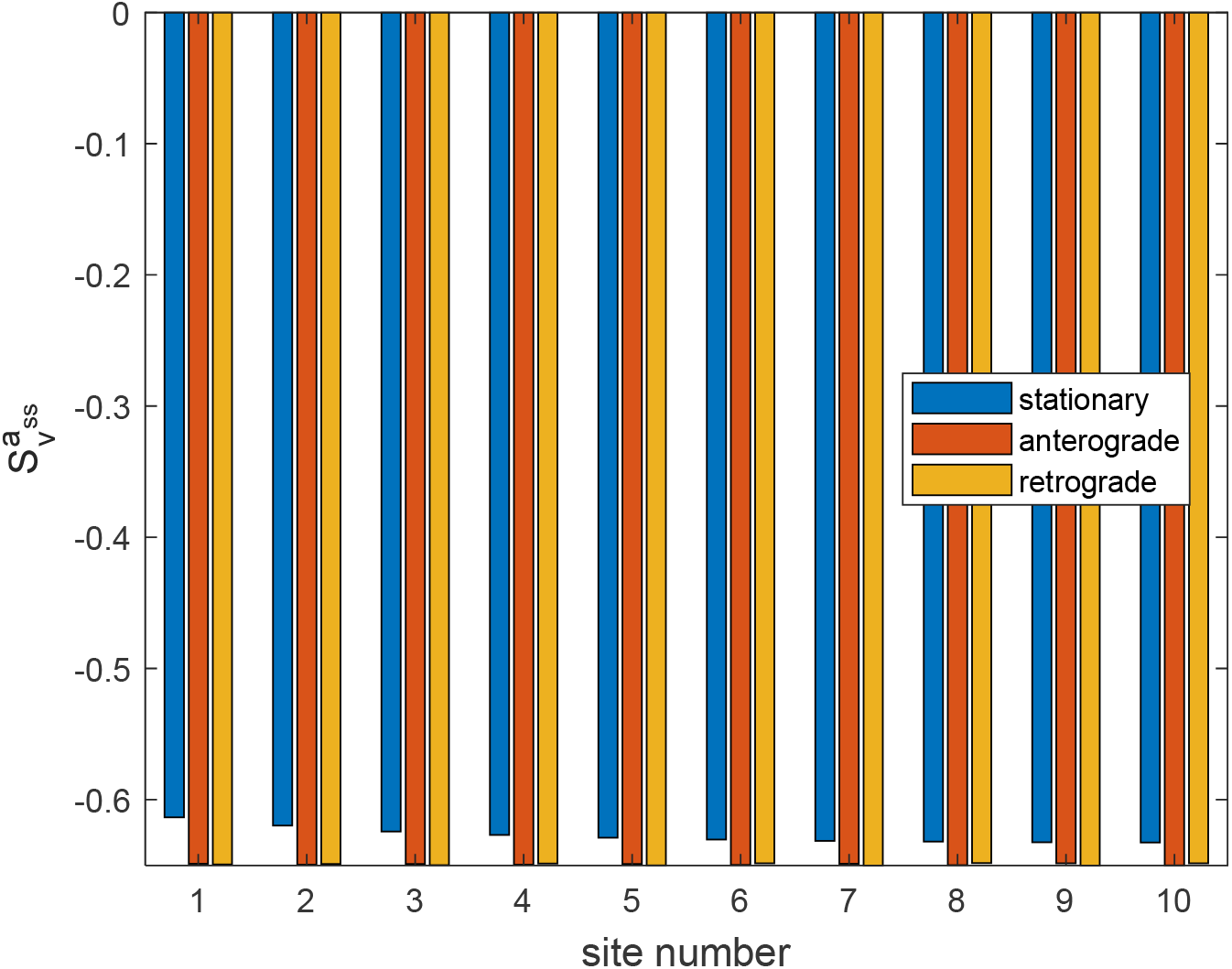
Dimensionless sensitivity of the mean age of mitochondria to the average velocity of anterogradely/retrogradely moving mitochondria, *v* (we assumed that *v_a_* = *v_r_* = *v*), versus the site number. Sensitivity is analyzed around *k_w_* = 5×10 ^4^ s^-1^, *p_s_* = 0.4. Computations were performed with Δ*v* =10^-1^ *v*. A close result was obtained for △*v* =10 ^2^*v*.

